# Self-supervised Benchmarking for scRNAseq Clustering

**DOI:** 10.1101/2023.07.07.548158

**Authors:** Scott R Tyler, Eric E Schadt, Ernesto Guccione

## Abstract

Interpretation of single cell RNAseq (scRNAseq) data are typically built upon clustering results and/or cell-cell topologies. However, the validation process is often exclusively left to bench biologists, which can take years and tens of thousands of dollars. Furthermore, a lack of objective ground-truth labels in complex biological datasets, has resulted in difficulties when benchmarking single cell analysis methods. Here, we address these gaps with count splitting, creating a cluster validation algorithm, accounting for Poisson sampling noise, and benchmark 120 pipelines using an independent test-set for ground-truth assessment, thus enabling the first *self-supervised* benchmark. Anti-correlation-based feature selection paired with locally weighted Louvain modularity on the Euclidean distance of 50 principal-components with cluster-validation showed the best performance of all tested pipelines for scRNAseq clustering, yielding reproducible biologically meaningful populations. These new approaches enabled the discovery of a novel metabolic gene signature associated with hepatocellular carcinoma survival time.

## Introduction

Identifying all biologically meaningful populations from large single cell datasets remains a challenge^1^. With current approaches to single cell RNAseq (scRNAseq) analyses, there are currently few options to validate clusters or topologies^2, 3^, none of which account for the fact that a cell’s position along a topology is partially attributable to the Poisson sampling noise, traceable to the random sample of the transcriptome that that was captured in the experiment. This lack of computational validation, accounting for sampling error, therefore, necessitates independent bench-validation of computationally inferred findings. When computational false discoveries are made, it can result in years of wasted time and large sums of money attempting to validate a non-existent population. False negatives on the other hand leave the scientific community blind to new potential discoveries.

Another large gap in single-cell methods benchmarking is the lack of objective “ground-truth” data to grade pipelines against each other in non-trivial datasets. While synthetic datasets provide a ground-truth, it is unclear how well they recapitulate all attributes of biologically derived data^4^. On the other hand, we cannot know *a priori* all attributes of biologically meaningful variation in complex biological data (i.e.: non-simulated), because these biological variations are what we seek to measure. Additionally, cluster labels by tissue experts are susceptible to human error and miss-classification of *unexpected* cell-identities^5^. In essence, an expectation, or hypothesis, can be a liability^6^. Furthermore, the primary goal of large scale - omics analyses is to develop *new* hypotheses agnostically of our prior expectations, to discover the unexpected, therefore leaving a large space for unsupervised analyses. However, unsupervised analyses are also not without their difficulties^1^.

A single round of clustering may not always be sufficient to uncover all biologically meaningful sub-populations in a dataset [i.e.: False Negatives (FNs)]. Conversely, a pipeline may yield false discovery of “too-many” clusters, splitting populations that only differ by random noise [i.e.: False Positives (FPs)]. While several heuristics can give users a sense for the structure delineated by cluster boundaries, others have recently shown that these may show inflated significance^2, 8^. Furthermore, it remains difficult to identify false-positives computationally, prior to the bench-validation attempt, because performing differentially expressed gene (DEG) analysis, or building a classifier to identify populations on the same data used to *generate* clusters/topology will nearly always give positive results, including false positives due to the “double-dipping” problem^5^.

In brief, this “double-dipping” problem is caused by the fact that a homogeneous population can be split in half (or any number of clusters), and the centroids of those two halves are separated by definition, simply because the population was over-clustered^8^. Therefore, when DEG analysis is performed, it is a self-fulfilling prophecy that there will be marker-DEGs, as these genes are what allowed these cells to occupy different regions of a Gaussian or uniform sphere in similarity space, even though they differ only by random sampling.

In Neufeld et al.’s recent work, they proposed a solution to the double-dipping problem in the form of count-splitting: in which each cell’s transcriptome is randomly segregated into two subsets that sum to the original observation^8^. These split counts can be used as a training- and test-set that can be treated as two independent observations of the same original cell, differing in which random slice of the transcriptome was observed by Poisson sampling. Therefore, if a topology is built for clustering, pseudotime, or other forms of analyses based on cell-cell similarity embeddings, the held-out split of the transcriptome can be used as an independent dataset with matched cells, thus preventing data leakage between topology building (on training data) and DEG or similar analyses assaying how gene expression changes along a topology (on test-data).

Here, we propose a within-cell 3-way split of single-cell RNAseq data, for initial topology building and clustering on the training-set, a validation-set for fine-tuning training results, and a test-set for DEG analysis. We created a graph-based cluster validation algorithm that uses this independent validation-split. Further, if we accept the premise that the primary purpose of clustering is to identify the greatest number of subsets that are reproducibly different from each- other, we can utilize DEG analysis on the test-set to independently confirm/refute cluster results. This enabled our creation of the first *self-supervised* benchmark in scRNAseq, quantifying performance of 120 clustering pipelines; this showed that anti-correlation-based feature selection, paired with our graph-based cluster validation notably increased overall performance across 37 benchmarking metrics. We also show that the top performing pipelines using our self- supervised benchmarking approach recapitulate previously described biology, and can lead to novel biological insights. Our work also shows a path towards other cluster/-topology-validation approaches in single-cell -omics, and towards the use of training-, validation-, and test-splits^9^ for future competitions, without reliance on consistency with manually defined cell-identity labels generated through use of prior authors’ preferred analysis pipelines^5^.

## Results

We first, created a new open source python-based bootstrapping implementation of the count-splitting method^8^, employing it to create three independent measures for training-, validation-, and test-sets that host each cell and gene in every split. Note that this differs from standard machine learning (ML) practice in which separate samples (cells in scRNAseq) would be used for training, validation, and test, looking for replicated characteristics in *different* populations. With count-splitting however, each cell has its own training-, validation-, and test- measure, allowing for labels discovered in training to be validated and tested for each observed cell (**Fig. 1a**).

**Figure 1:**
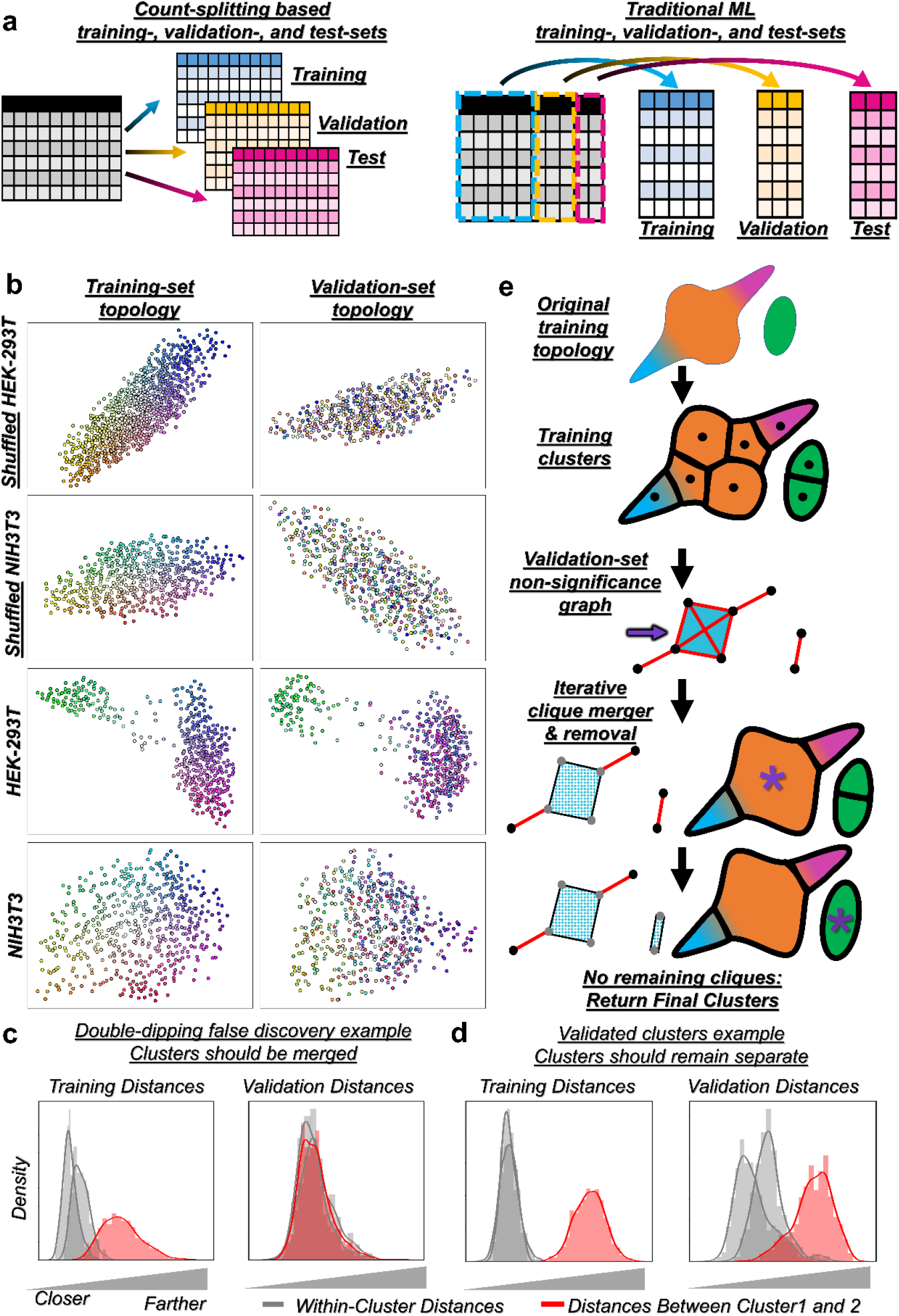
**Cluster-training and -validation premise. a**, Count-splitting training-, validation-, and test-sets provide random Poisson samples allocated to each split for every cell in a dataset, while traditional ML-style splits would randomly allocate different cells to training-, validation-, test-sets. **b**, kNN spring-embedding based topologies built from shuffled cell-line transcriptomes show global lack of topological concordance in the absence of biological signal; points correspond to cells, colorized based on training-data topology locations. **c**, An example histogram of within cluster distances (grey) which appear lower than between cluster distances (red), when using “double-dipped” training distances; however, the difference between the within vs between cluster distances did not replicate in the validation set, indicative of over- clustering. **d**, An example histogram in which clusters identified in the training-set replicated their separation in the validation-set. Histograms are colorized similarly to **c**. **e**, Schematic of cluster validation premise and iterative clique removal approach. Colored areas represent an observed topology upon which cells may lie; different colors represent biologically meaningfully distinct populations. Black bounding boxes represent cluster results. Validation-set distance comparisons (as in **b**,**c**) are used to connect clusters that are not significantly different from each other. Maximal cliques are identified and merged iteratively (purple stars), and removed from the network until no connected cliques remain, yielding the final cluster results.

We hypothesized that if a single *homogenous* population of cells exists, whose members differ only by Poisson sampling noise (i.e.: which random slice of the transcriptome was captured/observed), then if we examine two separate non-overlapping samples of the transcriptome, then these cells would be “re-randomized” in similarity space relative to the other cells. Therefore, this held-out validation slice would not show the same separation in similarity-/distance-space as those in the training-set used to define the clusters.

To test this hypothesis, we used *shuffled* transcriptomes of scRNAseq from two cell lines (guaranteeing no genuine structure), performing bootstrapped count-splitting to create two separate 50:50 transcriptomes for each cell, allocated at random. Indeed, if we examine the within and between cluster distances calculated in the training-set, these distances were not replicated in the validation set (**Fig. 1b; Supplemental Video 1,2**).

We assessed whether comparing within-cluster distances to between-cluster distances within the validation set may provide a statistical test for over-clustering. We applied this approach to the *unshuffled* versions of these cell-line transcriptomes where we expect a small, but biologically meaningful amount of variation. We compared the within-cluster and between-cluster distances, and indeed found some clusters where the *between*-cluster distances mirrored the *within*-cluster distances in the validation-set, indicating that over-clustering had occurred in the training-set (**Fig. 1b,c**), and that these clusters should be merged. Conversely, other cluster-cluster comparisons *preserved* the significant differences between these distributions, indicating that these clusters replicated, and should therefore remain separate (**Fig. 1b,d**; **Supplemental Videos 3,4**). In a complex dataset (mouse liver), global topology was highly conserved, yet some regions show discordance between training and validation splits, corresponding to Poisson sampling noise (**Supplemental Video 5**).

Importantly however, single-cell topologies may be continuous rather than discrete; yet it may still be useful to chunk the topology into bins (i.e.: clusters). We therefore created a cluster validation pipeline that can be combined with any processing and clustering pipeline, but uses the validation set to explicitly test whether the clusters found in training were recapitulated in the validation-set. This makes use of a statistical test comparing the within- to between- cluster distances in the validation-set, as described above, creating a weighted significance graph, in which clusters are connected to each other if they were not significantly different (**Fig. 1e**). We then iteratively merge graph cliques that are fully connected, based on the highest average edge p-value; in other words, if the clusters that are not significantly different from one another, iteratively merge the ones that are the *least* different on average. Note that this approach differs from other clustering approaches, some of which have built-in algorithms attempting to prevent over-clustering such as ChooseR^10^, Significance of Hierarchical Clustering (SHC)^11^, Phiclust^12^, due to our utilization of the independent within-cell validation-split.

To benchmark our approach, relative to the standard approach in which a single round of clustering is performed without validation, we created a recursive clustering pipeline that could mix-and-match feature selection algorithms, distance measures, and clustering algorithms. We further sought to assess the impact of the relative fraction of transcripts given to the training- or validation-sets (30, 40, 50, 60, and 70% of the transcriptome). These fully-crossed combinations resulted in 120 different analytic pipelines that we compared head-to-head.

We included the negative controls described above using *shuffled* cell-line scRNAseq, which must be called a single homogeneous population of cells; we also included the *unshuffled* versions of the cell-line data, in which only a single *round* of clustering should identify the subtle differences in cell-state, but should not be sub-clustered in an additional round of subdivisions (**Fig. 2a**). Passing negative controls however, while necessary for any scientific approach, is not sufficient however, as this does not yield information on the sensitivity to make true discoveries; we therefore also included more complex datasets, seeking to quantify the ability to make meaningful discoveries as well (**Fig. 2b**).

**Figure 2:**
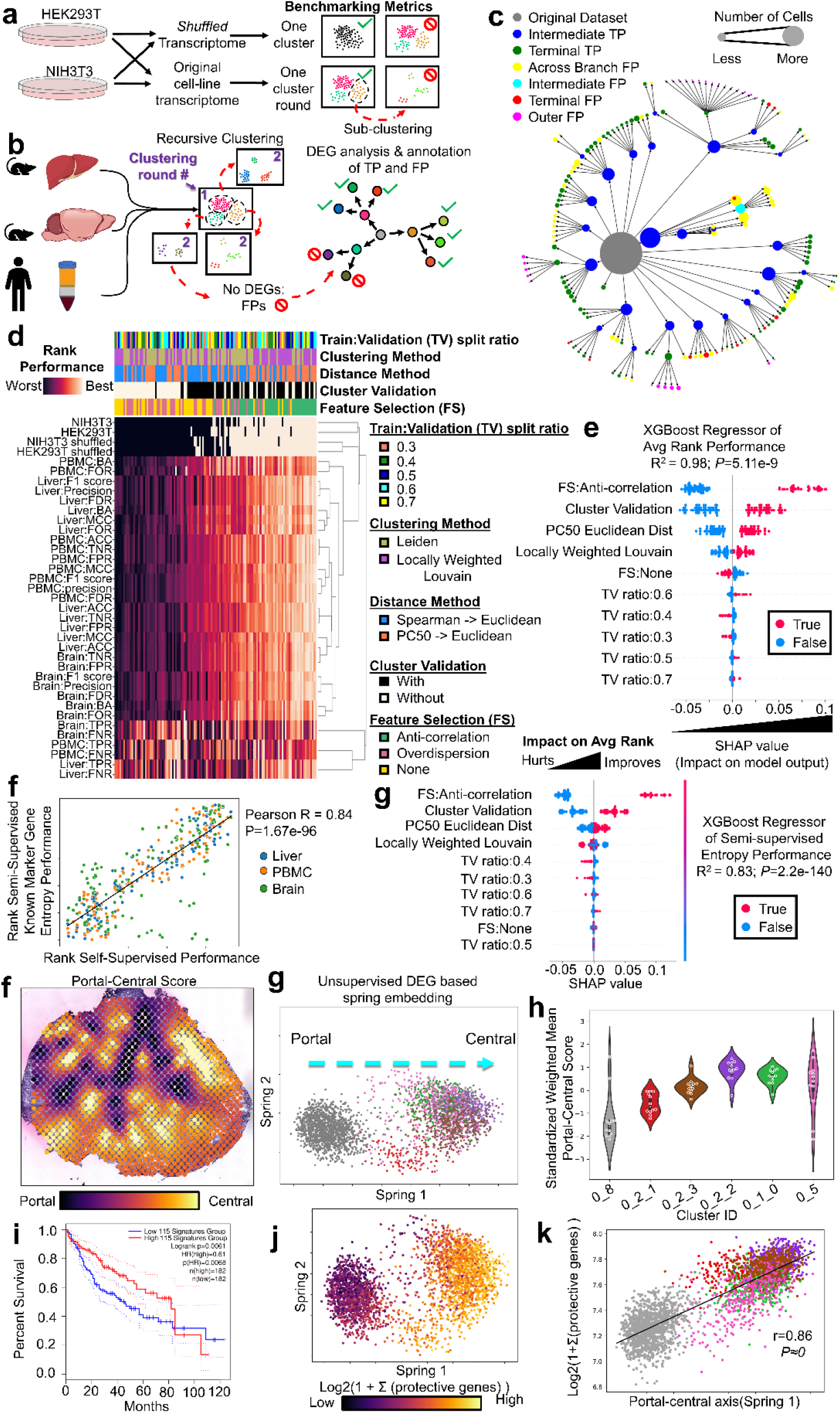
**Full benchmarking of cluster validation, using test-set based evaluation. a**, Benchmarking schematic with shuffled and unshuffled cell line transcriptomes, which should be identified as one cluster, or at most several clusters that should not be sub-clustered respectively. **b**, Benchmarking schematic using complex datasets. After recursive clustering, TP and FP subdivisions are classified based on presence/absence of DEGs in the test-set. These are then used to quantify terminal TPs (tTPs), intermediate TPs (iTPs), various forms of false positives (FPs), enabling calculation of relative FNs (rFNs) and relative TNs (rTNs), to create the full confusion matrix (See **Methods** and **ED** Fig. 1). **c**, An example of a recursive clustering tree, beginning with the original dataset (center-grey node), and progressively expanding outward for each successive round of clustering. Colors indicate TP/FP-type, and node size corresponds to the number of cells within the cluster. **d**, Heatmap showing ranking of all pipelines for all benchmark tasks; larger spots indicate better rank performance. **e**, Swarmplot of SHAP values quantifying the overall impact of each factor, relative to average performance, as predicted by the XGBoost regressor model of average rank performance in the benchmark. **f**, An example spatial transcriptomic dataset^15^, with portal-central scores assigned to each spatial voxel. **g**, Unsupervised spring-embedding based on differentially expressed genes between hepatocyte clusters. **h**, Violin plots showing standardized mean portal-central scores weighted by the relative deconvolved percentage of each spatial voxel for the shown clusters. This demonstrates that the clusters discovered by the top-performing pipeline identified the subtly variant spatial patterning of hepatocytes that correspond to the known portal-central axis. **i**, A Kaplan-Meier survival graph showing that expression of the combined significant gene signature are strongly associated with greater survival in HCC (*P*=6.1e-3, Log-rank p-value). **j**, The hepatocyte spring-embedding plot colorized by sum expression of the protective gene- signature. **k**, The sum expression of these genes are strongly correlated with the portal-central axis (Spring-1), colorized by cluster.

With complex biologically derived datasets however, we do not have a safe ground-truth assumption of either the correct number of clusters, or the cluster membership of individual cells. We began from the premise that the primary purpose of clustering is identification of the greatest number sub-divisions of a topology that are all *reproducibly* different from each-other. We can therefore use the *third* split of the data, to provide an independent measure of whether two populations indeed harbor statistical differences in expression. If two clusters do not harbor ≥10 DEGs up and down, these clusters were considered as false positives (FPs); conversely, if a cluster did harbor DEGs for all of its comparisons, this is considered a true positive (TP). We refine the TP population to identify the terminal TPs (tTPs), which demark each method’s most fine-grained clusters that were able to replicate in the test-set (see example recursive cluster graph in **Fig. 2c**, **ED Fig. 1**, See **Methods** for details). While computationally intense (2,312,006 cluster-cluster transcriptome-wide DEG analyses over the 120 pipelines compared), this approach allowed for an objective measure of apparent ground-truth without using prior expectations in the form of manual labels, or comparisons looking for *consistency* with the previously published labels generated by prior pipelines.

Despite having definitions for tTPs and FPs, it remained impossible to precisely define the number of negative populations (TN or FN), given that there is not a pre-defined set number of non-existent clusters. We can however, define *relative* negative metrics based on the maximum number of tTPs and FPs, taking these as maximum “total” number of “True” or “False” populations relative to the tested methods; this allowed for the calculation of relative TNs (rTNs) and FNs (rFNs) (**Fig. 2b,c**; See **Methods** for details). The calculation of these four metrics (tTP, FP, rTN, and rFN) opens the door to utilizing nearly all standard ML metrics for classifier efficacy, without user defined cell-type labels, which may be biased towards experimenter expectations, thus enabling a fully *self-supervised* benchmark.

With these four metrics in hand, we performed the full benchmark comparing 120 analysis pipelines, quantifying eleven metrics including: true positive rate (TPR), true negative rate (TNR), false positive rate (FPR), false negative rate (FNR), false discovery rate (FDR), false omission rate (FOR), precision, absolute accuracy (ACC), balanced accuracy (BA), Mathew’s correlation coefficient (MCC), and F1 score in each complex dataset, in addition to the pass/fail cell-line negative controls, summing to a total of 37 rankings for all 120 pipelines (**Fig. 2c**, **ED Fig. 2**, **ED Table 1a-c**).

Overall, we found that the pipeline with the best average rank was anticorrelation-based feature selection, followed by the Euclidean distance of the first 50 principal-components (PCs), paired with locally weighted Louvain modularity, and cluster-validation (60% training:40% validation) (**Fig. 2d**). To quantify the average individual impact of each variable [train:validation (TV) ratio, methods of feature selection, clustering, distance, and cluster validation], we built an XGboost regressor model for average rank performance, using these factors as explanatory variables. We then calculated the average impact of each variable on the predicted model output using Shapley (SHAP) values^13^. Fitting with the top performing pipeline, the largest contributors to average rank performance were 1) anti-correlation based feature selection, followed by 2) using cluster validation, then 3) calculating the Euclidean distance on the first 50 PCs (as opposed to Euclidean distance of cell-cell Spearman correlations), then 4) the use of *some* form of feature selection (either anti-correlation or overdispersion), while 5) the training:validation split ratio had comparatively little impact, but 0.6 training performed the best. (**Fig. 2e**).

A truly important aspect of unsupervised clustering is interpretability; in this context: “Did my pipeline discover biologically meaningful groups?” While unsupervised analyses can discover previously undescribed groups, they should *also* discover groups that track with previously described cell-types and their markers. We therefore sought an independent method of validating the biological interpretability of cluster results. To this end, we measured the mean relative entropy within clusters based on curated marker genes for each tissue^14^, similar to a previously described approach^7^; in essence, this provides a semi-supervised method to answer the question: “Within a given cluster, does this cluster show ‘order’ or ‘disorder’ based on its mean expression of known marker genes relative to shuffled cluster-labels?”. Indeed, there was an exceptionally strong correlation between our self-supervised benchmark’s rankings and the rankings based on cluster:marker-gene entropy (Pearson R=0.84, *P*=1.67e-96, **Fig. 2f**; **ED Table 1d**). Furthermore, building SHAP value explainers on an XGBoost regressor for marker gene entropy rank (**Fig. 2**), led to exceptionally concordant results with our semi-supervised benchmark’s results (**Fig. 2e**), with the top contributors to marker gene entropy performance being anti-correlated features selection, cluster validation, and PC50-Euclidean distance (**Fig. 2g**). This gives an orthogonal semi-supervised validation that our *self-supervised* benchmark not only incentivizes discovery of reproducible clusters, but that those clusters are biologically interpretable and meaningful.

To showcase this property, we examined whether the populations discovered by the top performant pipeline appeared to be biologically meaningful. To this end, we hypothesized that the mouse hepatocyte populations within the liver benchmark-dataset would show spatially-variant patterning via spatial transcriptomics^15, 16^ (**Fig. 2h,i**). We quantified the spatial variation scores for the top 50 marker genes for each hepatocyte population or 50 random genes using maxspin^16^; indeed, these top marker genes populations were significantly more spatially variant compared to a random sample of genes (**ED Fig. 2a,b**; T=3.83, *P*=5.0e-3). Furthermore, when quantifying a relative portal-central score for each cluster based on deconvoluted spatial abundance of each cluster, we found that these clusters were strikingly differentially abundant along the portal-central axis (**Fig. 2j**, F=13.9, *P*=2.68e-9, 1-way ANOVA, n=12 biological replicates). This finding matches well with the finding that an unsupervised spring-embedding showed concordant variation along the portal-central axis based on the expression of known marker genes of spatial zonation (**ED Fig. 2b**). Lastly, fully unsupervised DEG and pathway analysis showed the top significant pathways as relevant to changing metabolism moving from portal-to-central: amino-acid metabolism, arachidonic/fatty acid metabolism, metabolism of lipids, and PPAR/Wnt signaling (**ED Fig. 2c**), with additional clusters identified for proliferative hepatocytes cells. All of these findings are well supported within the literature including their spatial distributions^17^, thus confirming that the tTP clusters discovered through recursive clustering with the best performing pipeline identified by our self-supervised benchmark are biologically meaningful, even for populations of cells that only differ in their metabolic role.

Lastly, given that recent evidence has pointed towards metabolic reprogramming in hepatocellular carcinoma as an important aspect of cancer progression, we tested whether the gene expression programs differentially expressed between these tTP hepatocyte subsets were associated with cancer survival, after filtering out all genes associated with the cell cycle to prevent generic enrichment for proliferation, which would be expected to be negatively associated with survival. Indeed, survival was significantly associated with three of these gene- sets (**ED. Fig 3a,b**, *P*<0.05, Survival analysis, BH corrected Log-rank p-value). To our surprise however, each of these gene-sets appeared associated with *greater* survival time; we therefore combined these into a single combined protective gene signature in HCC, which was strongly associated with survival (**Fig. 2k**, **ED Table 2a**, *P*=6.1e-3, Log-rank p-value). Interestingly, expression of these genes were also strongly correlated with the portal-central axis (**Fig. 2l,m**, Spearman-rho=0.85, *P*≈0). Through pathway analysis, we found that the genes in these protective gene-sets were largely associated with metabolic pathways (**ED Table 2b**). This data fits with the fact that β-catenin is strongly associated with both central-hepatocyte cell identity and metabolic programming^18^, and that β-catenin driven HCC tumors grow more slowly^19^, yet are more recalcitrant to treatment^20^.

Overall, these results show proof-of-principle that the fully unsupervised approaches presented here, as discovered by our self-supervised benchmark, can yield robust biologically meaningful subsets, and lead to novel biological insights.

## Discussion

Here we present a count-splitting approach inspired by Neufeld et al^8^, that uses a training-set for topology building and clustering, a validation-set to optimize the training-clusters (or other topology-associated analysis), and a final test-set for DEG analysis. These splits enabled our creation of a new cluster validation algorithm and also allowed us to develop the first self-supervised benchmark of scRNAseq analysis pipelines. Our benchmarking not only uncovered a new approach to quantify state-of-the-art (SOTA) performance but also illuminated the most performant methods.

To make these approaches widely available and useful to the community, we have released open-source python packages of the bootstrapped count-splitting and our cluster validation algorithm, which python scanpy/AnnData framework^21^.

This test-set based approach to identify terminal true positives (tTPs) and false positives (FPs) allowed for additional calculation of relative TN and FN populations (rTN and rFN), completing the confusion matrix, granting access to the full suite of ML scoring metrics in a fully unsupervised manner. This approach also circumvents the critical limitation of prior benchmarks that necessitated a user to provide cluster labels – all of which were defined by the use of a prior publication’s preferred pipeline. This requirement therefore benchmarks for consistency with whatever methods were used by the prior publications, rather than benchmarking the ability to identify reproducibly different subsets of cells. We additionally observed concordance with our self-supervised benchmark ranking, and the ability of a pipeline’s clusters to yield clusters with high amounts of expression structure based on previously described marker genes. We further validated that the top performant pipeline as assessed by our new benchmarking method indeed identified biologically meaningful subsets through identification of the subtly metabolically different hepatocyte subsets, defined by spatial zonation patterns.

This self-supervised benchmark of 120 pipelines tested, using *100 billion* gene-level comparisons, ranking on the average of all 37 metrics quantified, show that anti-correlation based feature selection^22^ to be the greatest contributor, while the use of our cluster validation algorithm had second greatest impact on average performance, with a modest contribution also by performing Euclidean distance on the first 50 PCs as the most compared to Euclidean distances on Spearman correlations.

Moving forward, we anticipate that others will extend our general framework for continuous topology based analyses, such as RNA velocity^23, 24^. We further hope that this work will spur on the creation of other count-splitting based topology/cluster validation or uncertainty measures.

While the primary limitation of using a validation-split for clustering in addition to a test-split for DEG analysis is the loss of sensitivity, these limitations may be overcome through more recent bench-biology methods have dramatically increased efficiency of gene detection^25^.

Indeed, the PBMC, liver, and brain datasets used here were from third generation single-cell RNAseq using Parse Biosciences’ high-sensitivity V2 chemistry,^25^ which showed the ability to discover the sub-populations of hepatocytes that were spatially variant differing primarily in metabolic state, suggesting that sensitivity may not be a major limiting factor with high-sensitivity single-cell methods. Despite this however, the approaches that we employed will still be limited to identifying the populations that a given dataset has the power to detect through DEG analysis. Additionally, it is possible that two populations differ by linear or non-linear combinations of several genes, but none individually hold the power to cross a significance threshold. While non-DEG based methods could be employed in the test-set, we chose this approach because biological interpretations are frequently based on DEGs – however, there remains room for innovation in the exact methods used for identifying true and false discoveries with the test-set.

Overall, we have created the first count-splitting based cluster validation algorithm, that accounts for Poisson sampling noise, within the context of single-cell data analysis using count-splitting to create a training-, validation-, and test-set. We have also shown that cluster validation, along with anti-correlation based feature selection, are extremely effective at limiting computational false-discoveries while maintaining high true-discovery rates, validated by our confirmation of the spatial heterogeneity of these populations in the mouse liver. We also have shown that using the test-set to define a ground-truth can also enable unbiased *self-supervised* benchmarking, independent of experimenter prior expectations of which cell-types should be present, or which attributes of variation should be labeled as biologically meaningful. We hope that these methods will save bench biologists from the frequent task of attempting to validate computational false-discoveries mediated by overfitting the training data; we also hope this work inspires new computational validation approaches in single cell analyses and benchmarks.

## Acknowledgements

Support for this work was provided by K99HG011270.

## Author Contributions

SRT conceived of and performed all analysis and wrote manuscript. EES and EG advised and edited manuscript.

## Competing Interests Statement

Authors declare no conflicts of interest.

## Methods

### Code and Data Availability

The code for count splitting is available at the repository: https://bitbucket.org/scottyler892/count_split/src/main/

And is also pip installable: python3 -m pip install count_split

The code for this benchmark are located at the repository below: https://bitbucket.org/scottyler892/count_split_clust_bench/src/master/

All data used for benchmarking is available in the data.zip file distributed in the repository.

The clustering validation package is available here: https://github.com/scottyler89/dclustval

https://dclustval.readthedocs.io/en/latest/index.html

and is pip installable:

python3 -m pip install dclustval

#### Cell line data

NIH3T3 cells and HEK293T cell data was previously released from 10x genomics and analyzed^22^. The used splits are stored in AnnData format^21^ in the data.zip file.

#### Shuffled cell-line data

Within-gene bootstrap shuffling was performed to ensure that there was no biologically meaningful signal remaining within the data, while still being biologically derived. These are also included in the data.zip file within the benchmarking repository.

#### Example cell line topologies shown

In **Fig. 1b-e**, we show several example topologies and within/between-cluster distances. These analyses were performed using a popular approach with Scanpy’s overdispersion based feature selection algorithm, followed by PCA reduction to 50 principal components, followed by generation of a k-nearest neighbor embedding (described later in the methods in greater detail). The example topologies shown are a spring embedding of a kNN (described in detail in a later section) built from scanpy’s implementation of overdispersion based feature selection, followed by PCA reduction to the first 50 PCs, followed by the cell-cell Euclidean distance calculation.

This pipeline for the example display was selected due to the broad use of these methods. The code for this process is located within the bin/plot_diff_between_train_val_topologies.py file within the benchmarking pipeline.

#### Cluster train-, validation-, and test-splits

For all benchmarking measures, the test-split was 35% of total counts in the original dataset. The 30, 40, 50, 60, and 70% splits noted in the manuscript are related to the percentage of the split remaining after the 35% test-set was removed. The final ratios of train, validation, test are:

30: train:19.5, validation:45.5, test:35 40: train:26, validation:39, test:35

50: train:32.5, validation:32.5, test:35 60: train:39, validation:26, test:35

70: train:45.5, validation:19.5, test:35

Splits were made using the bin/run_splits.py file in the benchmarking repository.

#### Feature selection algorithms

Anti-correlation-based feature selection was implemented using default parameters from the anticor_features package with the get_anti_cor_genes function, with defaults used for other parameters using the log(1+counts) matrix.

Overdispersion-based feature selection was performed as implemented in scanpy^21^ with the author suggested parameters as follows:

sc.pp.normalize_total(adata, target_sum=1e4) sc.pp.log1p(adata)

sc.pp.highly_variable_genes(adata, min_mean=0.0125, max_mean=3, min_disp=0.5)

#### Distance algorithms

Either the Euclidean distance of the cell-cell Spearman correlation matrix was taken, or the Euclidean distance of the first 50 principal components on the log transformed data was taken. Euclidean distances were calculated using the euclidean_distances function from the sklearn package^26^, spearman correlations were calculated by the no_p_spear function in the anticor_features package^22^.

#### Clustering algorithms

Locally weighted Louvain modularity was implemented using default parameters from the bio- pyminer package with the do_louvain_primary_clustering function^27^.

Leiden modularity was implemented with default hyperparameters from the leidenalg package with the following call: leidenalg.find_partition(G, leidenalg.ModularityVertexPartition)^28^.

The k nearest neighbor graph, G, used for the Leiden modularity algorithm was defined with k as the maximum of 10 or the natural log of the number of cells. Neighbors were defined as the k cells with the lowest values from whichever distance matrix was calculated as described above.

#### Recursive clustering

Recursive clustering results are modeled as a directed graph network, using the networkx package in python^29^. The representation of the dataset begins as a single node representing the full dataset, with neighbors being defined as the clusters identified from clustering the cells within the parent node. Each subsequent neighbor was recursively sub- clustered, and represented as neighbors in the directed graph, until a given pipeline found only a single cluster.

#### Cluster validation algorithm

When the cluster validation algorithm was employed, it was applied at each round of recursive clustering based on the distances in the validation set. The exact same pipeline of feature selection and distance measures was applied to both the training and validation sets separately. This yields two distance matrices, the training distances (TD) and validation distances (VD). The clusters identified by the training algorithm were then compared pairwise based on the validation distances. Notably, the fact that the full processing pipeline was applied to the two splits independently means that the noise, variability, or error introduced at each processing step is propagated forward.

Within each round of clustering, each cluster-cluster pair (as determined by the given clustering algorithm on the training distances), the within-cluster distances were subset from the distance matrix:

With a vector (L) of length N(cells), containing the cluster labels, that may range from 1 through N(clusters), a list of lists CL is created of length m to catalogue the indices for each cluster- label.

For w=1..N(clusters), and i=1..N(cells):

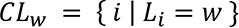

Next, a vector of each cluster’s within-cluster distance (WCD) is subset from the validation distance matrix (VD). For w=1..m (number of clusters) and i=1..n and j=1..n, where n is the number of cells in the dataset.

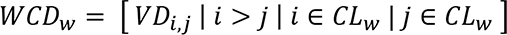

Next between-cluster distances (BCD) were subset for all cluster-cluster pairs: For w=1..m and x=1..m where m is the number of clusters.

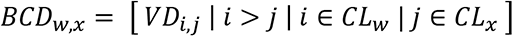

Note that the *i* > *j* condition is simply to subset the upper-triangle of the distances, thus preventing double-counting.

Next, we perform a statistical test to quantify, for each cluster-cluster pair, whether the BCD distances are significantly greater than the WCD for each cluster. The test applied is a simple t- test of independence on the rank transformed data using the ttest_ind function in scipy^30^. These p-values are stored in a significance matrix S of dimension m by m where m is the number of clusters.

For w=1..m and x=1..m clusters:

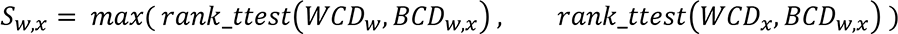

Next, these p-values within S are adjusted using the Benjamini-Hochberg (BH) FDR correction.

S is then masked for any values less than an alpha significance threshold (0.01 by default). For w=1..m and x=1..m clusters:

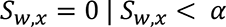

Next, S is used as a weighted adjacency matrix to generate an undirected graph representation of “non-significance,” G. In this graph, vertices represent the original clusters discovered in training, and included edges are those representing a cluster-cluster comparison that was non- significant. Weights correspond to the adjusted p-values described above contained within the significance matrix S.

Next, we enter an iterative loop within which, the maximal clique(s) is/are identified via the networkx find_cliques function^29^. If there exists a tie for the maximal clique (based on number of vertexes), the clique with the highest average edge weight is selected. Because the weights correspond to adjusted p-value, this is the maximal clique that has the highest average adjusted p-value is selected, or in other words the least significantly different maximal clique.

This clique is then removed from the graph, and the cells that corresponded to the clusters within this clique are combined into a single cluster within the final labels (FL) for that round of clustering. Note that removal of a vertex within the maximal clique may alter or disrupt the presence or size of a different clique that it was previously a member of; this means that this iterative maximal clique removal process must be carried out sequentially to determine the number of final clusters.

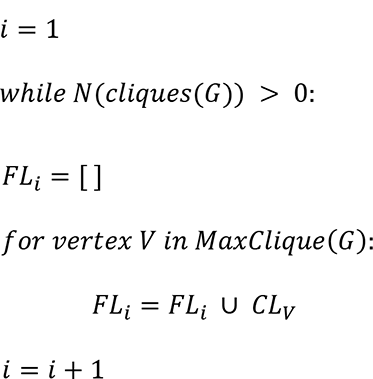

This maximal clique removal procedure is repeated iteratively until no more cliques are remaining within the graph, and any orphan vertices remain their own cluster, thus yielding the final cluster labels.

*for vertex V in orphans(G):*

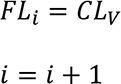

#### Differential expression analysis on recursive clustering results

Differential expression was performed for all pairwise clusters that were neighbors of the same parent node. Prior to DEG analysis, for expediency, the data was subset to include only genes that were expressed in at least five cells within either cluster. Differential expression was only ever performed on the third test-split of the data, which was not used in any way prior to DEG analysis. DE was performed by a linear model on dense rank transformed expression, using a binary encoding vector for cluster membership using the OLS function in the python statsmodels package^31^. For each cluster-cluster transcriptome-wide comparison, the p-values were then FDR adjusted via the BH correction as implemented in the statsmodels function: statsmodels.stats.multitest import fdrcorrection^31^.

#### Definition of TP, tTP, iTP, and FP clusters

With the differential expression as defined above, for each cluster, we calculated the minimum number of differentially expressed genes between each cluster and its other neighbors relative to the shared parent node within the recursive cluster graph. Differentially expressed genes that were up or down regulated were quantified separately. We classified a TP as a cluster that had ≥10 significantly up, and ≥10 down genes relative to each of its cluster comparisons. It is important to quantify both up and down genes in this classification due to the effect of depth or capture efficacy. For example, a single biological population could have its transcriptome sampled at depths ranging an order of magnitude or more in single cell datasets. Sub-dividing this population into high-counts and low-counts populations would yield apparent differential expression simply due to depth effects, even after counts per 10k normalization. At the same time, it has been long understood that biologically distinct populations harbor different amounts of RNA^32, 33^, meaning that using total counts as a covariate in the model could regress out the biologically meaningful signal because total counts would be confounded with this biological signal. To circumvent these two problems, we use the simple metric of requiring both ≥10 up, and ≥10 down regulated significant differences in the test-set in all cluster comparisons to be considered a TP cluster. Any cluster that does not have at least 10 significantly up and down genes compared to the other cluster is considered a FP. The alpha threshold for significance was 0.001 applied to the BH corrected p-values (**ED Fig. 1a**).

For TPs, we aimed to further identify each pipelines’ maximum number of informative clusters that replicated via DEG analysis in the test-set. However, two different algorithms may come to the same final cluster results in two different ways. For example, one algorithm may find 16 final clusters in a single round of clustering, while another algorithm may find the same 16 final clusters through iterative bifurcations. The first algorithm will have a total of 16 cluster vertexes in the total recursive cluster graph. However, the second algorithm would have a total of 2 in the first round of clustering, 4 nodes in the second, 8 in the third, and 16 in the fourth, yielding a total of 30 total nodes in the recursion graph, all of which may harbor DEGs.

However, the same 16 clusters exist as the most fine grained TPs between these two approaches.

We therefore required a refined TP count that quantifies specifically the *terminal* TPs (tTPs) that replicated by DEG in the test-set; under this scenario, the bifurcating graph and single-shot results that landed upon the same final clusters would be rewarded equally. To this end, the algorithm begins at the leaves of the recursive clustering tree, and traverses inward until it reaches the first instance of a TP node; this node is labeled as a terminal TP (tTP). The remaining TPs upstream of the terminal TPs are labeled as intermediate TPs (iTPs) (**ED Fig. 1b**). Next, the minimal connected graph that contains the tTPs (called a Steiner tree) is identified using the networkx steiner_tree algorithm^29^.

In some cases, there were also FPs at an intermediate stage, upstream of a tTP, these are labeled as intermediate FPs (iFPs). Terminal FPs are FPs that exist in the same branch and cluster level as a tTP. Lastly, all truly informative subdivisions of a dataset should differ from one-another, even if they were sub-clustered across different branches of the clustering tree.

Notably however, inaccuracies in clustering at round 1 may result in an inflated number of within-branch tTPs, that if compared across branches would be uncovered as across-branch FPs (**ED Fig. 1c**). To this end, as a final check for cluster utility and redundancy, we performed the above described DEG analysis comparing all within branch tTPs to each other across all branches of the graph, therefore converting some of the within branch tTPs to across branch FPs. Those within branch tTPs that remained different from all others were retained within the final tTPs.

#### Calculation of relative true negatives (rTN) and relative false negatives (rFN)

While the use of the test-set allowed for quantification of terminal true positives (tTP) and the various forms of false positives (FP) including iFPs, tFPs, and outer FPs, this approach did not explicitly enable the calculation of true or false *negative* populations. This is because there is a near infinite number of ways to define errant subdivisions of a dataset. However, TN and FN are critical measures for the calculation of many traditional machine learning (ML) classification metrics. We therefore sought to calculate *relative* TN and FN lying on the scale of what was discovered by all the assessed pipelines for a given dataset.

We begin from the premise that the perfect clustering method would yield the maximum subdivisions of the dataset that, such that for each subdivision, there would be reproducible differences between itself and all other subdivisions; in other words, tTPs as defined above.

This would be a maximally sensitive clustering algorithm, that also does not create erroneous subdivisions that are not significantly different from other subdivisions.

The calculation of relative false negatives was derived from the maximum number of TP found by any algorithm for each dataset independently. Discovery of fewer true positive populations is taken to indicate lesser sensitivity. To this end, with tTP representing the vector corresponding to each pipeline’s number of tTPs:

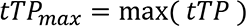

Therefore the relative false negatives for each pipeline *i* is:

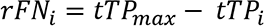

For *i*=1..n, where n is the number of pipelines tested, and rFN corresponding to the vector of rFN values for each pipeline *i*.

Unlike calculating rFN, calculation of an absolute number of true negatives, requires knowledge of all possible wrong cluster results. However, there is a gargantuan number of all possible permutations of cell-cell pairings of clusters of differing sizes and combinations. We therefore again sought a relative true negative (rTN) metric. Because each pipeline will find its own version of reproducible or non-reproducible clusters, these populations are not necessarily zero sum; we therefore cannot simply calculate the maximum total number of clusters found for each pipeline then subtract the tTP, FP, and rFN populations to yield the rTN population. To illustrate this point, we entertain hypothetical results in which, the pipeline with greatest sensitivity discovered 60 tTPs, and the pipeline giving the greatest number of total clusters, found 65 clusters (tTP+FP), here called C_max_.

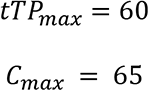

For a given method that found 10 tTPs, and 25 FPs, the calculation of rFN is relatively simple:

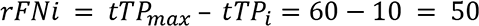

However, summing tTPi, FPi, and rFNi yields a value greater than Cmax:

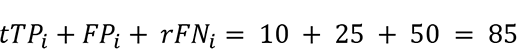

We therefore, instead, imagine a scenario in which the total number of positives is equal to the highest number of tTPs of all pipelines, and the highest number of possible negatives being the highest number of FPs (iFP, tFP, outer FP, or across-branch FP) all classified as FPs. This yields a maximum number of total clusters (TC_max_) to be:

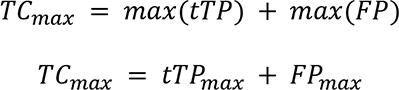

This allows for the zero-sum, calculation of the number of relative true negatives (rTN) as follows:

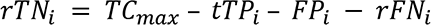

However, because: *rFN*_*i*_ = *TP*_*max*_ − *tTP*_*i*_, we can expand *rFN*_*i*_ as follows:

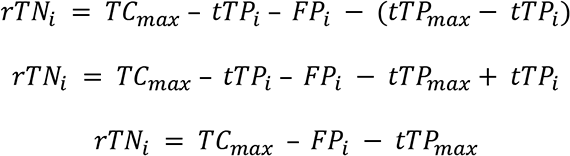

Similarly, because *TC*_*max*_ = *tTP*_*max*_ + *FP*_*max*_ we can expanded *TC*_*max*_ as follows:

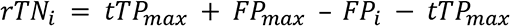

Which reduces to:

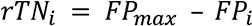

Thus we have the relative true negatives (rTN) are simply the maximum number of false positives within the dataset, minus the number of false positives for the given pipeline *i*, which as shown above respects the theoretical perfect confusion matrix in which the total positives are tTPmax, and total negatives are FPmax, with the total possible clusters as previously defined (*TC*_*max*_ = *tTP*_*max*_ + *FP*_*max*_); yet this formulation of rTN and rFN also allows for dynamically defined clusters for each pipeline, so long as we can calculate tTP and FP.

#### Marker gene entropy analysis

We first calculated each cluster’s mean expression of the given tissue’s marker genes (as defined by CellMarker2.0^14^). The collated subset we used is also available in our repository in the data/manual_markers_cellmarker2.0.tsv file in our repository. First the genes, then the clusters were sum normalized such that the sum values of each cluster’s mean expression vector totaled 1. This matrix was then used to calculate normalized entropy, which is calculated via the scipy.stats.entropy function divided by the log of the number of features, which is bounded between 0 and 1 invariantly to the number of features (in this case, genes) used.

Because each tissue had different marker genes, and cannot be directly compared between tissues, the normalized entropy’s were rank transformed and linear normalized within tissues for the final rank entropy metric. These analyses are contained in the bin/do_entropy_analysis.py file within the primary benchmarking repository.

#### QC of hepatocyte dataset

As an example to show the biological relevance of tTP clusters identified by the top performing pipeline, we examined the hepatocytes of the liver dataset used in the benchmark (included in repository). The percentage of total counts corresponding to mitochondrial encoded genes was calculated by summing the counts of genes leading with the sub-string “mt-”.

Interestingly, we found that the cluster with highest mitochondrial percentage (cluster 0_9_0) also showed very high levels of pseudo-genes (as calculated by selecting genes beginning with “Gm[0-9]+” used as a regular expression) (**ED Fig. 2a**). This may be attributable to the fact that many pseudogenes correspond to the ancient capture of the proto-mitochondrial bacteria, which resulted in 612 nuclear insertions of the mitochondrial genome, which have since evolved into pseudogenes^34^. This may therefore manifest as mis-mapping of mitochondrial transcripts to these nuclear encoded pseudogenes.

In connected regions of this topology, were two other clusters (0_0 and 0_10) with low total counts, and high percentage of counts mapped to nuclear-LncRNA, indicative of the nucleus containing fragments that may have ripped off from the anucleated, mitochondria high fragments. Therefore, 0_9_0, 0_0, and 0_10 were removed for subsequent analyses.

#### Spatial transcriptomic analysis of liver

Spatial transcriptomic datasets of mouse liver were available in h5ad format through a prior publication^15^. Using the top 50 marker genes (ranked by mean expression difference between the highest expressing hepatocyte cluster and second highest) from each tTP hepatocyte population (defined as cluster mean expression Z-score >0 for expression of albumin), were assayed for spatial information with the python package maxspin, which calculates spatial mutual information relative to shuffled version of the dataset^16^. A 1-sample T-test was performed to compare the percentage of these genes that were significantly spatially variant as determined by maxspin compared to the percentage of a random set of 50 genes using the scipy ttest_ind function. This analysis is shown in **ED. Fig. 2a**.

To quantify each spatial puncta for the relative abundance of each cluster, deconvolution was performed using RNAStereoscope^35^ as implemented in scvi-tools^36^. We sought to use these deconvolutions to quantify whether the relative distribution of hepatocyte clusters across the liver were related to the portal-central axis. To this end we first calculated a portal-central score and used these along with the deconvolved relative abundances to identify relative spatial variation along the portal-central axis as described below.

Twelve biological replicates were included within the spatial transcriptomic datasets in the prior publication^15^. Contained within the distributed .h5af files were distances to the nearest central and portal veins (*Distcentral* and *Distportal* respecively)^15^. We used these distances to calculate a single metric to capture the portal-central axis (*PCscore*), with the following formula:

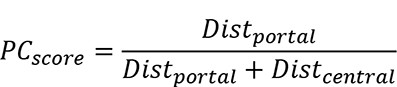

This gives a score bounded between zero and one that corresponds to which type of vein the spatial voxel was closer to. Given this formula, if a spatial voxel was located very close to a portal vein, then it would have a low relative value, whereas a high value would correspond to being very close to a central vein. An example of this, overlayed on H&E histology is shown in **Fig. 2f**.

Here, each voxel has a deconvolved relative abundance of each cell type, as well as a *PCscore*; we sought to use each of these metrics to quantify the spatial variation of each cluster along the portal-central axis. To this end, we modified the deconvolution relative abundances to quantify the relative distribution of each cluster across all voxels. That is – within a single cluster, quantifying each voxel’s contribution to the total abundance of that cluster.

Given a (v x c) deconvolution matrix (**D**) with v voxels and c clusters, and an output within-cluster relative abundance matrix (**A**):

For i=1..*N*(v) voxels and j=1..*N*(c) clusters:

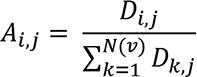

Such that all the sum of all columns (corresponding to clusters) equals 1. To calculate a weighted average cluster-level portal-central score (CPC_score_), we then applied the following formula for each slide:

For j=1..N(c) clusters:

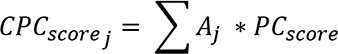

With *A*_*j*_ corresponding to an N(voxel) length vector containing the relative abundance of a cluster for all voxels (as defined above), and *PC*_*score*_ corresponding to an N(voxel) length vector for the portal-central scores described above. This gives a single portal-central score for each cluster, for each slide. Lastly, to control for slide-specific effects, each slide’s results (*CPC*_*score*_) were standardized (mean-centered and divided by the standard deviation).

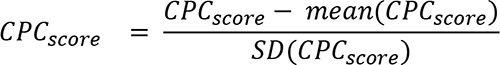

These standardized scores are displayed in **Fig. 2h**.

All of the above described analyses are contained within the “graph_deep_dive.py” script contained within the bin directory of the main benchmarking repository: https://bitbucket.org/scottyler892/count_split_clust_bench/src/master/.

#### Hepatocyte spring-embedding

The spring-embedding of hepatocytes was based off of the expression of the subset of genes that showed differential expression (as defined above) between any of the pairwise comparisons between clusters. While these DEGs were identified on the test split, the display was calculated on the training split, which is designated for topology building. Fitting with the best-practices identified within the self-supervised benchmark, a kNN was built (as described above) using the Euclidean distance on the first 50 PCs from this DEG subset. Positions were then calculated using the networkx spring_layout function on the kNN graph.

#### Survival analysis on hepatocyte sub-cluster DEGs

The DEG lists comparing hepatocytes (methods described above), were first filtered to remove cell-cycle associated genes. This was done to prevent the generic and uninteresting finding that proliferation associated genes result in worse survival; this was particularly important because of the presence of the highly proliferative cluster (0_5). This was done by first querying gProfiler for the full gene list mapping to the cell-cycle GO term (GO:0007049)^37^. The remaining non-cell-cycle related significantly differential expressed genes were then split into two lists for each comparison, those that were higher in one cluster, and those that were higher in the other. These lists were used for survival analysis using the gepia python package^38^, which references TCGA data^39^, using the sv=gepia.survival() function. Given that we were examining hepatocytes, this analysis was limited to liver hepatocellular carcinoma (HCC), abbreviated (LIHC) in gepia. This is done by setting the dataset parameter as follows: sv.setParam(key=’dataset’,value=[“LIHC”]). Log rank p-values were Benjamini-Hochberg corrected using the scipy function fdrcorrection. The gene-signature of the expression of these genes was calculated by taking the log2 sum of the expression of all genes in the gene list (**ED Table 2a**). The correlation between this sum expression and the portal-central axis (Spring-1 coordinates) was assessed by the scipy pearsonr function. Pathway analysis was performed by via the gProfiler python package, using a custom background comprised of genes that were detectably expressed within the test split (from which DEGs were derived)^37^.

These analyses are contained within the “graph_deep_dive.py” script contained within the bin directory of the main benchmarking repository.

---

**Extended Data Figure 1:**
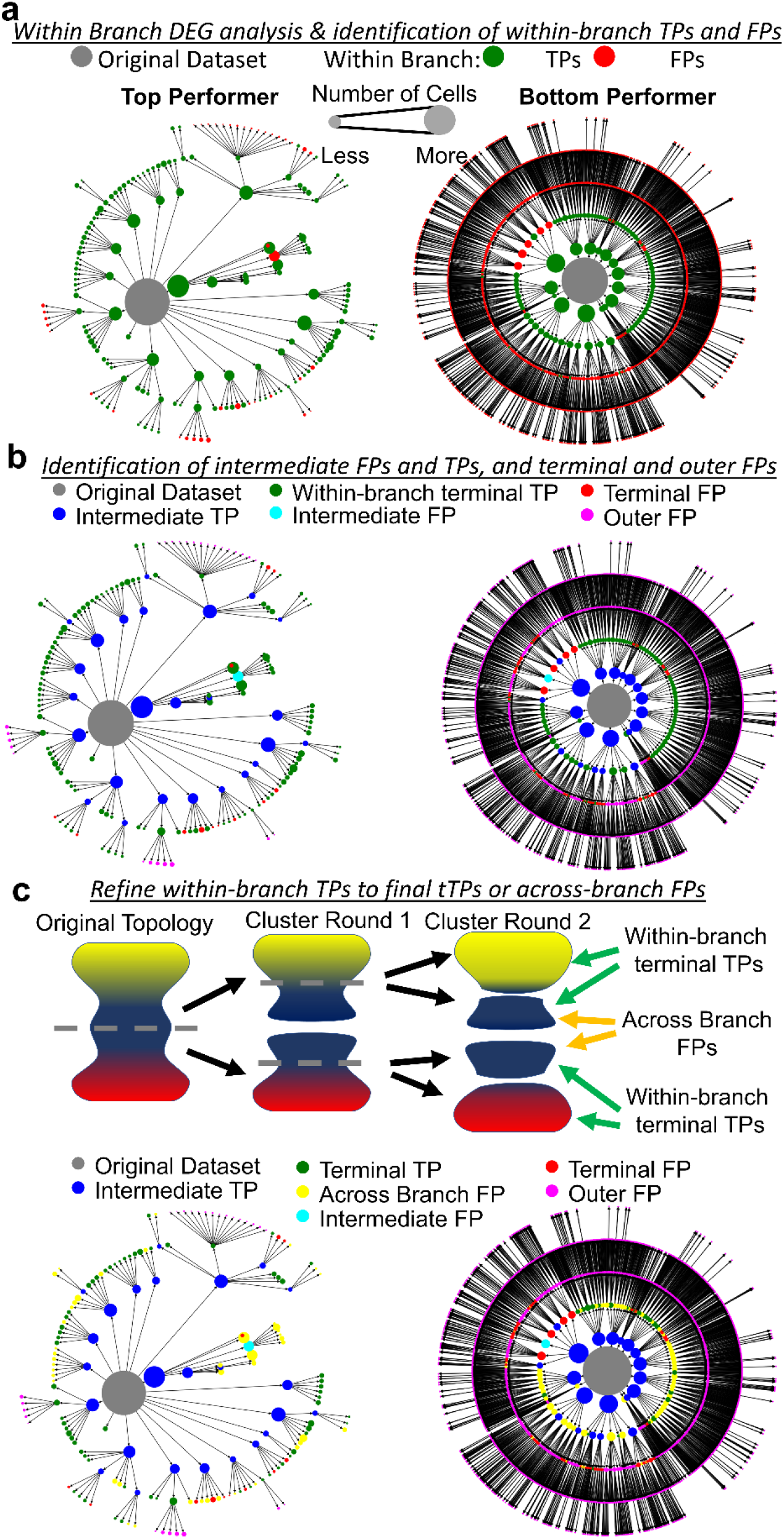
**Annotation of TP, FP, tTP, iTP, rFN, rTN, and tFP clusters using the test-set. a**, Example recursive clustering graphs from the top and bottom performing pipelines, colorized by the initial round of DEG analysis to identify within-branch and within- recursion level FP and TP populations. **b**, Within-branch terminal TP subset (tTP) are identified by first subsetting the graph to contain the minimal tree containing all of the TPs (identified within **a**) (also called a Steiner Tree). The terminal leaves of this subgraph are the tTPs, with other TPs internal to this subgraph are called intermediate TPs (iTPs). Within-branch tTPs therefore quantify each branch’s most fine-grained replicated clusters, while iTPs are simply subdivisions along the way towards this greatest level of granular subdivision. Intermediate FPs (iFPs) were also identified within these graphs. Terminal FPs (tFPs) are clusters at the same recursion level as a given branch’s tTPs, outer FPs are errant subdivisions that were not contained within the TP-containing Steiner tree. **c**, A final potential failure mode exists within the tTPs, where a single group may have been split upstream in the recursion tree, creating the illusion of two separate tTPs. These are identified by performing DEG analysis between all of the tTPs identified in **b**; if any tTP-tTP comparisons were not significantly different, they were relabeled as across-branch FPs, as this indicates an errant split upstream in the clustering tree, while those groups which showed significant differences between all comparisons were maintained as final tTPs.

**Extended Data Figure 2:**
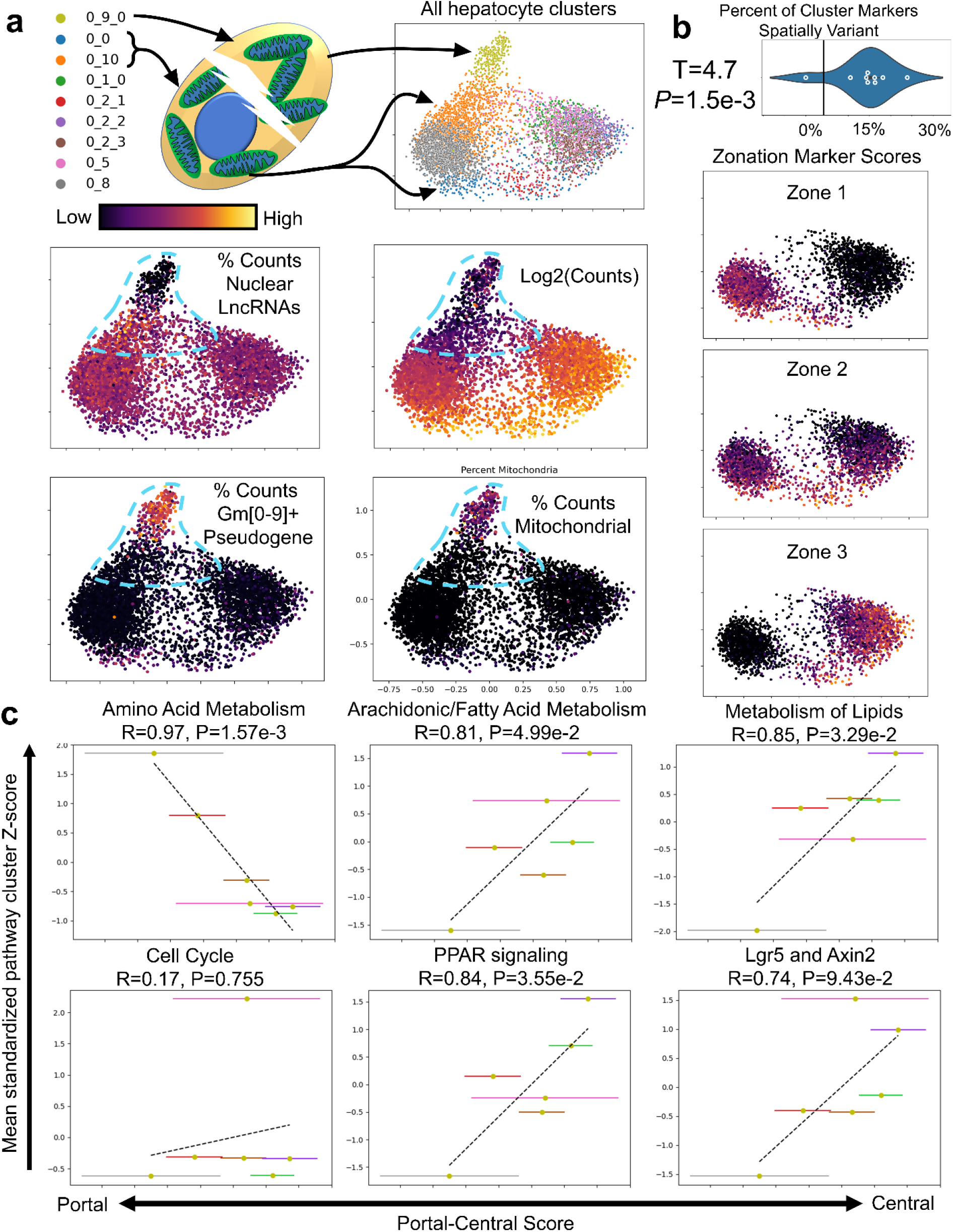
**Hepatocyte sub-clusters**. **a**, Unsupervised spring embeddings based on differentially expressed genes between hepatocyte clusters are shown, colorized by various quality control (QC) metrics. These QC metrics identify cell fragments included in the dataset, highlighted in blue dotted outline. Cluster 0_9 showed high levels of mitochondrial and pseudogene content with no detected nuclear long non-coding RNAs (lncRNAs) (Xist, Malat1, and Neat1), while the connected clusters 0_0, and 0_10 showed low total counts, and modestly higher than typical nuclear lncRNA content. These clusters were removed for subsequent analysis. **b**, A violin-plot compares the top-50 marker genes for each cluster, compared to 50 random genes (black line), for the percentage that are spatially variant as determined by maxspin^16^, using previously published liver spatial transcriptomic datasets^15^. Significantly more of the cluster marker genes were significantly spatially variant compared to randomly selected genes (T=4.7, *P*=1.5e-3, 1-sample T-test). Plotting known liver zonation marker scores (color) on unsupervised spring-embeddings confirm that clusters track with known spatial zonation markers. **c**, Clusters are metabolically variant in a manner that correlates with previously described as varying with zonation^15, 40^.

**Extended Data Figure 3:**
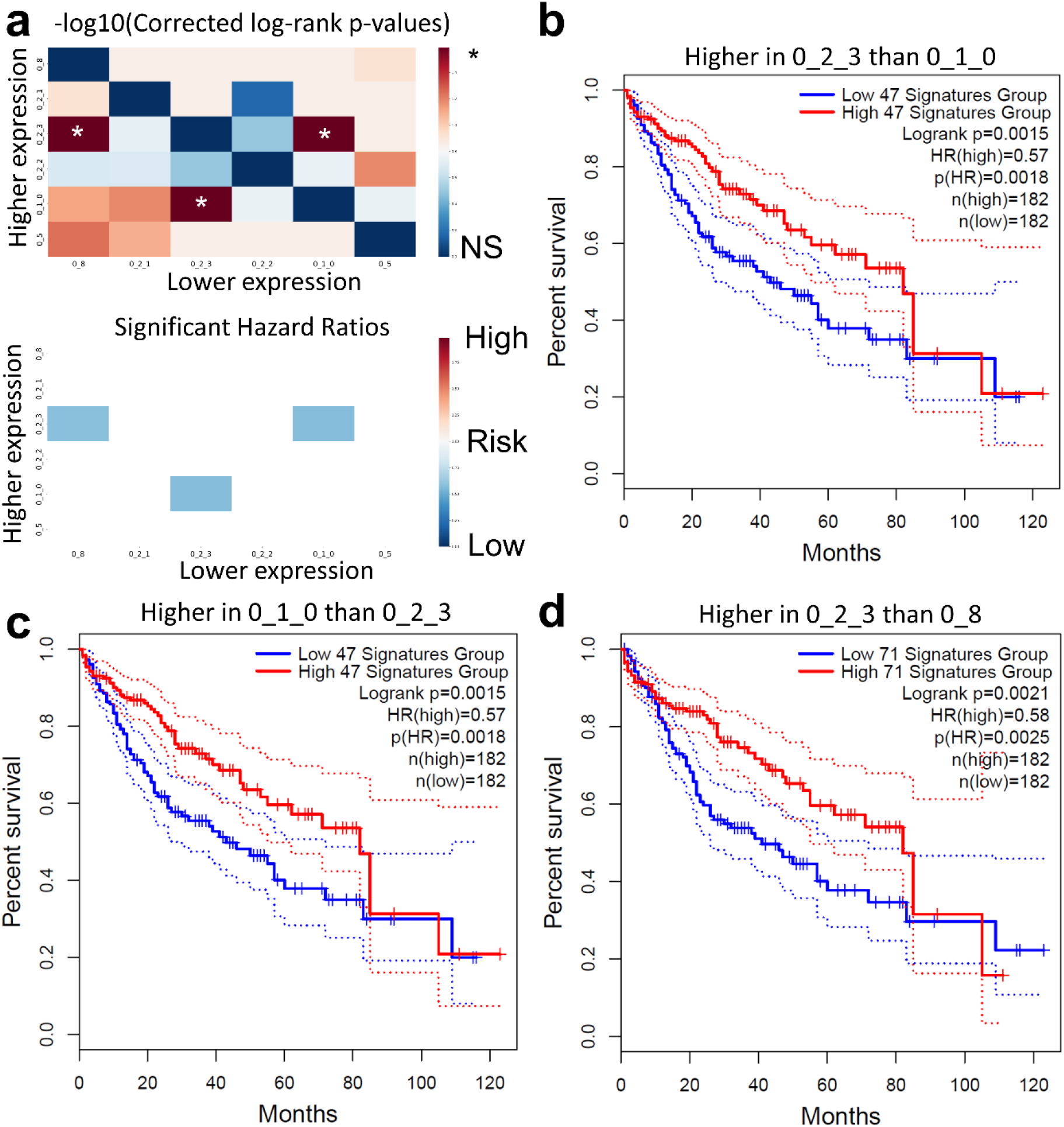
**Hepatocyte gene signatures associated with hepatocellular carcinoma survival. a**, A heatmap of log-rank significance for the pairwise DEG lists in their association with HCC survival. Also displayed is a heatmap of the hazard ratio, masked to only display significant results. **b-d**, The individual Kaplan-Meier survival plots for each of the three significant signatures. Note that despite the similar output between these survival curves, these were indeed differing gene-lists, but likely stratified patients similarly or equivalently. Nominal p-values are shown within the Kaplan-Meier plots, while those shown in panel **a** are Benjamini- Hochberg corrected.

**Extended Data Table 1: Benchmark results. a**, The cell-line based metrics used for all benchmarking results for the 120 pipelines tested. **b**, The metrics used for benchmarking the 120 pipelines tested that were based on complex biological datasets. **c**, The sorted rank results of the benchmark.

**Extended Data Table 2: Hepatocellular carcinoma protective gene signature. a**, The list of genes that were contained within the combined protective gene signatures. **b**, The pathways associated with the genes associated survival.

**Supplemental Video 1: Lack of concordance in topologies between training and validation sets when no structure exists in a dataset.** Animation of shuffled HEK293T cell transcriptomes with spring-embedding based locations from the training-set (beginning of video), and their corresponding validation-set topology locations (end of video). Cells are colorized based on their original coordinates within the training-set topology throughout the video.

**Supplemental Video 2: Lack of concordance in topologies between training and validation sets when no structure exists in shuffled NIH3T3 transcriptomes.** Animation of shuffled NIH3T3 transcriptomes with spring-embedding based locations from the training-set (beginning of video), and their corresponding validation-set topology locations (end of video). Cells are colorized based on their original coordinates within the training-set topology throughout the video.

**Supplemental Video 3: Mixture of concordant and discordant topologies when minimal dataset structure exists.** Animation of the original (unshuffled) HEK293T cell transcriptomes with spring-embedding based locations from the training-set (beginning of video), and their corresponding validation-set topology locations (end of video). Cells are colorized based on their original coordinates within the training-set topology throughout the video. The left-right domains remained separated in both training and validation sets, while the cells within the right domain were scrambled along the vertical axis, indicating that this elongated sub-topology was not replicated.

**Supplemental Video 4: Mixture of concordant and discordant topologies when minimal dataset structure exists.** Animation of the unshuffled NIH3T3 cell transcriptomes with spring-embedding based locations from the training-set (beginning of video), and their corresponding validation-set topology locations (end of video). Cells are colorized based on their original coordinates within the training-set topology throughout the video. Cells tended to stay within their left-right domains, while the up-down axis was less conserved in this system.

**Supplemental Video 5: Example of variation in training and validation topology a complex biological dataset**. A spring embedding from a liver dataset shows overall conservation of global structure, while some regions show discordance, particularly with the region at the top of the plot in green show what appears to be cluster splitting and re-mergers between training and validation sets, representing a non-reproducible region that differs only by Poisson sampling noise.

## Notes

https://bitbucket.org/scottyler892/count_split/src/main/

https://bitbucket.org/scottyler892/count_split_clust_bench/src/master/

https://github.com/scottyler89/dclustval

https://dclustval.readthedocs.io/en/latest/index.html

### Competing Interest Statement

The authors have declared no competing interest.

